# Using expert driven machine learning to enhance dynamic metabolomics data analysis

**DOI:** 10.1101/482224

**Authors:** Charlie Beirnaert, Laura Peeters, Pieter Meysman, Wout Bittremieux, Kenn Foubert, Deborah Custers, Anastasia Van der Auwera, Matthias Cuykx, Luc Pieters, Adrian Covaci, Kris Laukens

## Abstract

Data analysis for metabolomics is undergoing rapid progress thanks to the proliferation of novel tools and the standardization of existing workflows. However, as datasets and experiments continue to increase in size and complexity, standardized workflows are often not sufficient. In addition, as the ground truth for metabolomics experiments is intrinsically unknown, there is no way to critically evaluate the performance of tools. Here, we investigate the problem of dynamic multi-class metabolomics experiments using a simulated dataset with a known ground truth and evaluate the performance of tinderesting, a new and intuitive tool based on gathering expert knowledge to be used in machine learning, and compare it to EDGE, a statistical method for sequence data. This paper presents three novel outcomes. First we present a way to simulate dynamic metabolomics data with a known ground truth based on ordinary differential equations. This method is made available through the MetaboLouise R package. Second, we show that the EDGE tool, originally developed for genomics data analysis, is highly performant in analyzing dynamic case vs control metabolomics data. Last, we introduce the tinderesting method to analyse more complex dynamic metabolomics experiments that performs on par with statistical methods. This tool consists of a Shiny app for collecting expert knowledge, which in turn is used to train a machine learning model to emulate the decision process of the expert. This approach does not replace traditional data analysis workflows for metabolomics, but can provide additional information, improved performance or easier interpretation of results. The advantage is that the tool is agnostic to the complexity of the experiment, and thus is easier to use in advanced setups. All code for the presented analysis, MetaboLouise and tinderesting are freely available.

## Introduction

The field of metabolomics, which studies small molecules inside organisms, has seen rapid growth over the last two decades and the amount of data being generated by metabolomics experiments keeps increasing. Although for many “typical” experiments standardized workflows are available, for more complex experiments such as dynamic metabolomics, sometimes also called longitudinal or time-resolved metabolomics, standardized workflows are lacking. With experiments gaining complexity, manual data processing and analysis becomes less feasible.

Over the last years many initiatives have focused on providing free and open access workflows and pipelines for metabolomics data analysis. For example, Workflow4Metabolomics [1] (W4M) provides an intuitive way of constructing workflows by linking modules together. These modules provide a multitude of steps for preprocessing, statistics, normalization and others. Many of these tools were originally written in R, python, etc. but have been converted to the Galaxy [2] environment that underlies W4M. The net result is that anyone now has access to the most advanced tools, but more importantly the workflows can be automated, standardized and shared, thereby greatly improving verifiability and reproducibility of research and results. This push towards reproducibility is, for example, also apparent in the new release of MetaboAnalyst [3], which is accompanied by the R package MetaboAnalystR [4] to allow researchers to easily share their workflow code for identifying compounds.

A specific type of metabolomics research consists of dynamic metabolomics experiments. Small molecules and their processes are the direct result of biochemical activity and therefore, metabolomics describes an inherently dynamic process. There is a large variety of dynamic metabolomics experimental setups, for example studying specific hormone levels in a single patient over a single day to find recurring patterns relating to biorhythm, or quantifying the treatment effect for a certain known metabolite, etc. An overview with examples and approaches to analyze these experiments can be found in Smilde et al. [5]. This paper focuses on the data analysis of untargeted dynamic metabolomics experiments. We can illustrate the need for such experiments by considering the use case of prodrug metabolism. Aspirin (acetylsalicylic acid), for example, is a prodrug and as such is not pharmacologically active. When taking this medication the body effectively starts converting it into the active medicine. In this process certain metabolites are consumed and others are formed over a period of time. If it is unknown which metabolites are formed from possible active compounds or prodrugs, untargeted experiments can deliver answers. However, the data processing of such dynamic experiments is not trivial, as there are often multiple sample classes measured over multiple time points.

There is no common default strategy yet for the data analysis of large-scale, untargeted, dynamic metabolomics experiments in which, simultaneously, a large number of features need to be considered but also a large variety of different time profiles. One promising approach mentioned by Smilde et al. [5] is EDGE [6, 7]. This tool originates from the genomics field and uses natural cubic splines. Even though it is not widely used in metabolomics it provides a promising solution for discriminating different time profiles. The drawback is that EDGE is specifically tailored to two-class problems. Using EDGE for multi-class experiments is not straightforward as problems arise both on the multiple testing aspect and on the combination of individual two-class comparisons (which is further elaborated in the Materials and Methods section).

In this paper we propose a workflow that provides an easy and intuitive solution to these problems. This is done by incorporating expert knowledge into the analysis pipeline, in a playful manner that we implemented in an app called “tinderesting”. Experts are iteratively shown figures of data entities, for example features, and rate each based on whether the feature is interesting or not. Next, this labeled data provided by the experts is used to train a machine learning model that in turn can be used to analyze the full dataset. The idea is in spirit analogous to the recaptcha concept, in which internet users are required to decipher jumbled-up words that are presented on the screen, assisting in elucidating hard to digitize text from old manuscripts [9]. The suggested method inherently tries to emulate the decision process of the expert. Note that this also means that if the expert is biased, the model will be biased as wel. Specific precautions need to be taken to avoid such biases, as discussed in the Discussion & Conclusion section. The advantage of our approach is that it is simple to perform and, in comparison with EDGE, it is agnostic to the number of sample classes present in the data. The one prerequisite is that the data can be readily visualized in a way that offers the expert a quick view on whether a feature is interesting or not. This is indeed the case in dynamic metabolomics experiments.

## Materials and methods

### Validation dataset: dynamic metabolomics data

To demonstrate EDGE and tinderesting we first work with a simulated dataset. The advantage of this approach is that the dataset can be constructed in such a way that the ground truth is known. However, constructing such a simulated longitudinal metabolomics dataset is not common practice. We propose a dedicated method for simulating these data, based on a limited number of biologically inspired assumptions. We compare the resulting dataset to an actual dynamic metabolomics dataset to evaluate to which extent the simulated data is realistic.

#### Simulating dynamic metabolomics data

To simulate longitudinal metabolomics data we will use the following biologically inspired assumptions:

1. the dynamics in the data are governed by an underlying network with an appropriate connectivity distribution (the Baabási-Albert model is chosen [10, 11], see Supporting Information S1 for further discussion),
2. nodes in the network are metabolites, which have certain starting concentrations that evolve over time,
3. transitions in concentrations are caused by enzymes. Effectively these are the rates/fluxes that govern flow in the network,
4. some of these enzymes are assigned to multiple edges and are also influenced by the adjacent nodes/metabolites. That way, an external influx in metabolite X can cause a depletion of metabolite Y somewhere else in the network (X increases, thus rate/flux of X to X’ increases, this is the same rate/flux as the one from Y to Y’, so Y depletes),
5. the intake of certain compounds (e.g. nutrients) causes an external influx in some metabolites, (temporary rise in concentration),
6. the rate/flux increases or decreases depending on the concentration of the metabolite causing the reaction. This increasing rate is limited and follows a sigmoidal curve.

To construct the dataset twenty different networks are generated, consisting of 50 metabolites (nodes) each. To these networks one or both of the following are added: enzymes (rates) or external influxes. The different sample classes are differentiated by what is added to the network:

- sample (with influx and includes enzymes),
- blank (no influx, includes enzymes),
- negative control (with influx, no enzymes).

Each sample has three replicates, constructed by initializing the network with a constant concentration with a random Gaussian term. Each sample is ‘measured’ at eight different time points. To obtain a dataset with a ground truth, there will be only one of twenty networks that receives an influx (sample and negative control). In the other networks there is no influx. This construction results in 50 metabolites that are affected because they are in a network that undergoes changes originating from external influxes. These and only these are the metabolites of interest. Effectively this results in a ground truth where 5% of the 1000 metabolites are biologically relevant. These are the metabolites that the tools should label as significant. This setup is visualized in Fig 1.

**Fig 1.**
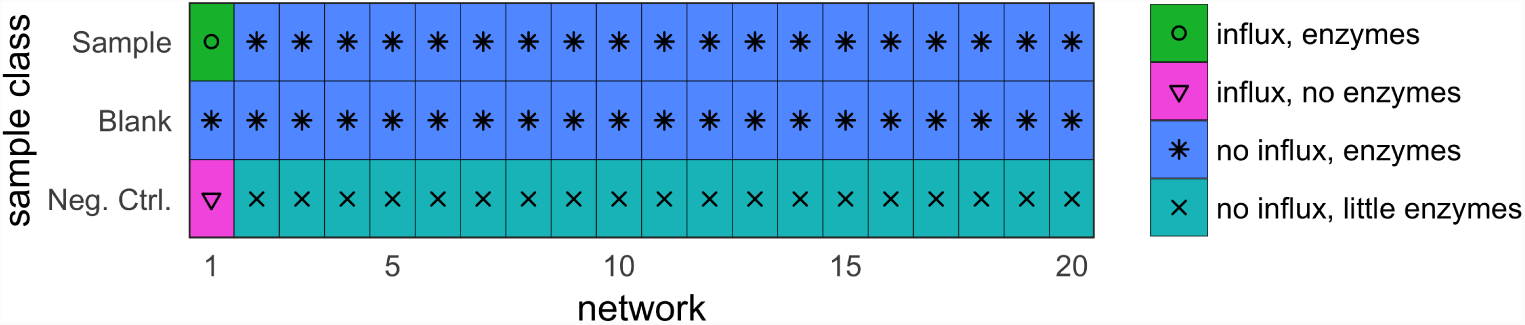
Visualized simulation setup. Only in the first network there are metabolites that differ between sample and blank as well as between sample and negative control. For the remaining 95% of metabolites there is only a small difference in level of enzymes between sample and negative control. These enzymes convert one metabolite into the other with a rate dependent on the enzyme quantity. For the sample vs blank case the only difference is the omnipresent noise on starting concentrations.

The concentration flow in the network is governed by ordinary differential equations [12] that are solved numerically by using the standard Euler equations with a sufficiently small time step. More specific information can be found in the Supporting Information S2 along with a small example network to illustrate the mathematics of changes in the network over time. The parameters for simulating these networks are discussed as well as how to construct networks that have a connectivity distribution according to a power law [13]. All code, examples, and instructions how to perform these simulations can be found in the MetaboLouise R package on github.com/Beirnaert/MetaboLouise and CRAN (awaiting approval).

#### Statistical methods for differential metabolism

The EDGE method [6, 7], originating from genomics, is used to find the features that exhibit differential metabolism over time. The main differences between differential expression over time of genes and metabolites are the scaling and transformations that are applied to the data. Differential gene expression data is often log transformed because roughly half the data points are in the [0,1] range, with 1 signifying no difference in expression, and the other half in the range [1, *∞* [. By log transforming these data the distribution changes to half of the data below 0 and half above 0. For liquid chromatography-mass spectromatry (LC-MS) or nuclear magnetic resonance (NMR) spectroscopy metabolomics data this problem does not occur. We are working with intensities for which all data are in the range [0, *∞* [. Therefore we apply a linear transformation so that each feature has a maximal intensity of 100 (divide by the maximum and multiply by 100).

EDGE works in two steps to compare whether two groups exhibit significant differential expression over time [6]. In the first step the null hypothesis is formed stating that there is no difference between the two population time curves. A single natural cubic spline curve is fitted to all the data and the goodness of fit is calculated. Splines are chosen to keep the method flexible enough to capture different types of dynamics. In the second step, the alternative hypothesis is formed and a separate curve is fitted to each group (also a cubic spline). See Fig 2 for a visualization of these two steps. A likelihood ratio test is performed to obtain a p-value that quantifies the improvement in goodness of fit. A large improvement signifies more evidence for a difference in time profiles between classes. Note that p-value corrections such as false discovery rate (FDR) controlling procedures are required in the case of multiple testing, see Supporting Information S3 for further details regarding multiple testing correction and EDGE [14].

**Fig 2.**
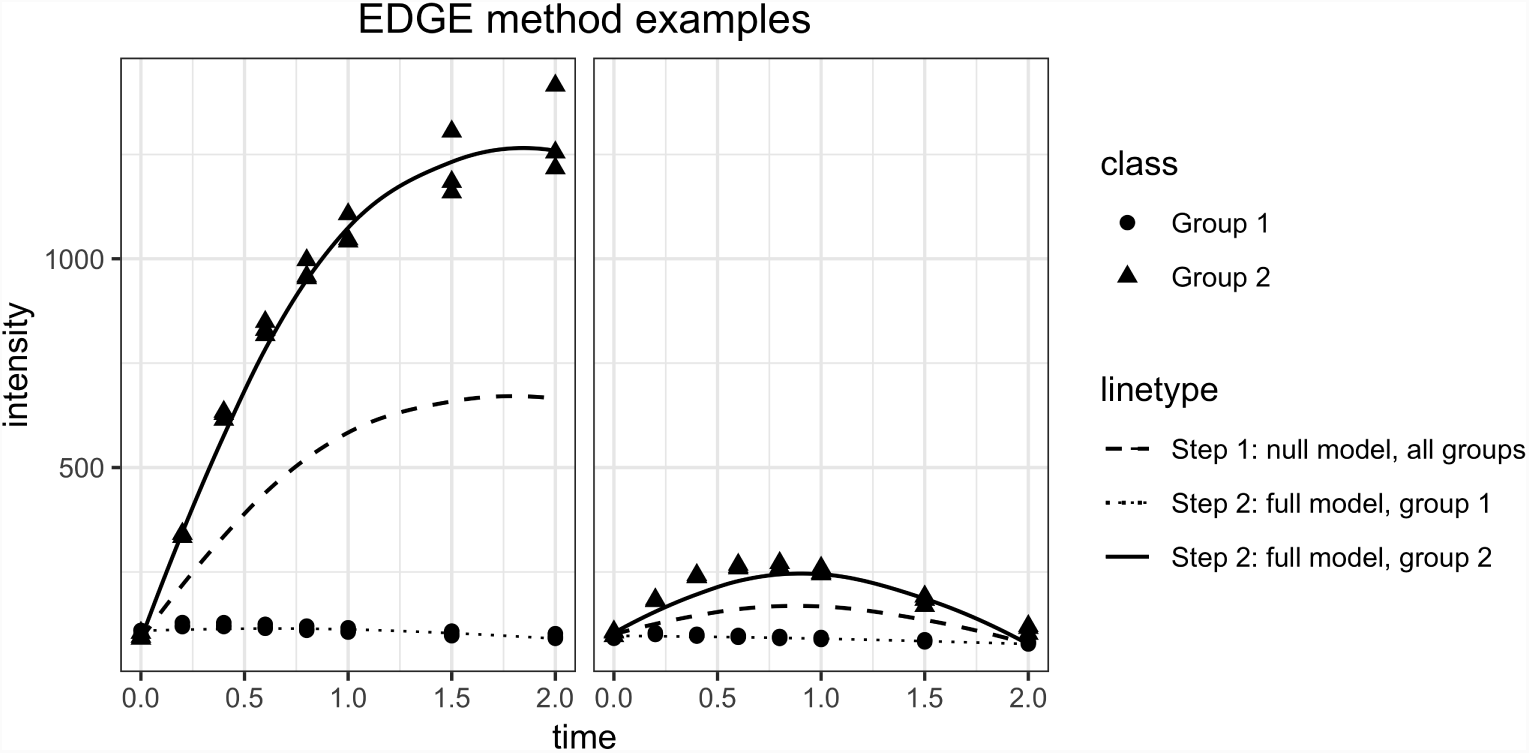
Illustration of the two steps in the EDGE algorithm for 2 different features (left, right). Step 1: the null hypothesis states that there is no difference between the groups, a single cubic spline curve is fitted to all data (dashed line). Step 2: in the alternative hypothesis a curve is fitted to each group separately (dotted and full line). The improvement in goodness of fit is a measure for the difference between the groups.

### Machine learning and tinderesting

The aim is to use a machine learning (ML) model that emulates the human revision process. The reason for this is twofold. First, the end result of statistical analysis generally contains false positives. Instead of manually looking at all the results to filter out these false positives, the machine learning model can be trained to do so. Second, in case of multiple biological sample classes in a single experiment, it can become non-trivial to interpret or combine the results of statistical methods. To train the ML model, labeled data is needed. These labeled data come from the tinderesting Shiny app, see Supporting Information S4 for an image of the full app, which is used by the experts to review the features. The experts have three options for labelling a feature: interesting, uninteresting or unknown. These correspond to the tinderesting labels. The unknown label indicates edge cases where the expert is uncertain. Features with the unknown label are omitted when constructing the final random forest model. Thus, the final model is built by using the features that have a tinderesting label that is interesting or not. These features serve as the samples for the machine learning model (every sample to classify corresponds to a metabolic feature rated by tinderesting) and the machine learning features are the individual elements of those metabolic features i.e. an intensity for each sample class, replicate and time point combination. The performance is evaluated by receiver operator characteristic (ROC) with accompanying area under the curve (AUC) values as well as by precision-recall (PR) curves which provide results that are less influenced by class imbalances. Cross validation (10 fold) is used with 20 repeats (samples redivided over folds) to get an estimate of the variability of the ROC and PR curves [15]. In this paper, the default parameters are used for building the random forest model, see Supporting information S5 for the values of these parameters and performance comparisons to a naive Bayes classifier and support vector machine. Note that depending on the data, it can be beneficial to tune these parameters. After training the machine learning model, it can be used to score new data or rescore the entire dataset to look for potential false positives or false negatives in earlier statistical results. See Fig 3 for a visual overview.

**Fig 3.**
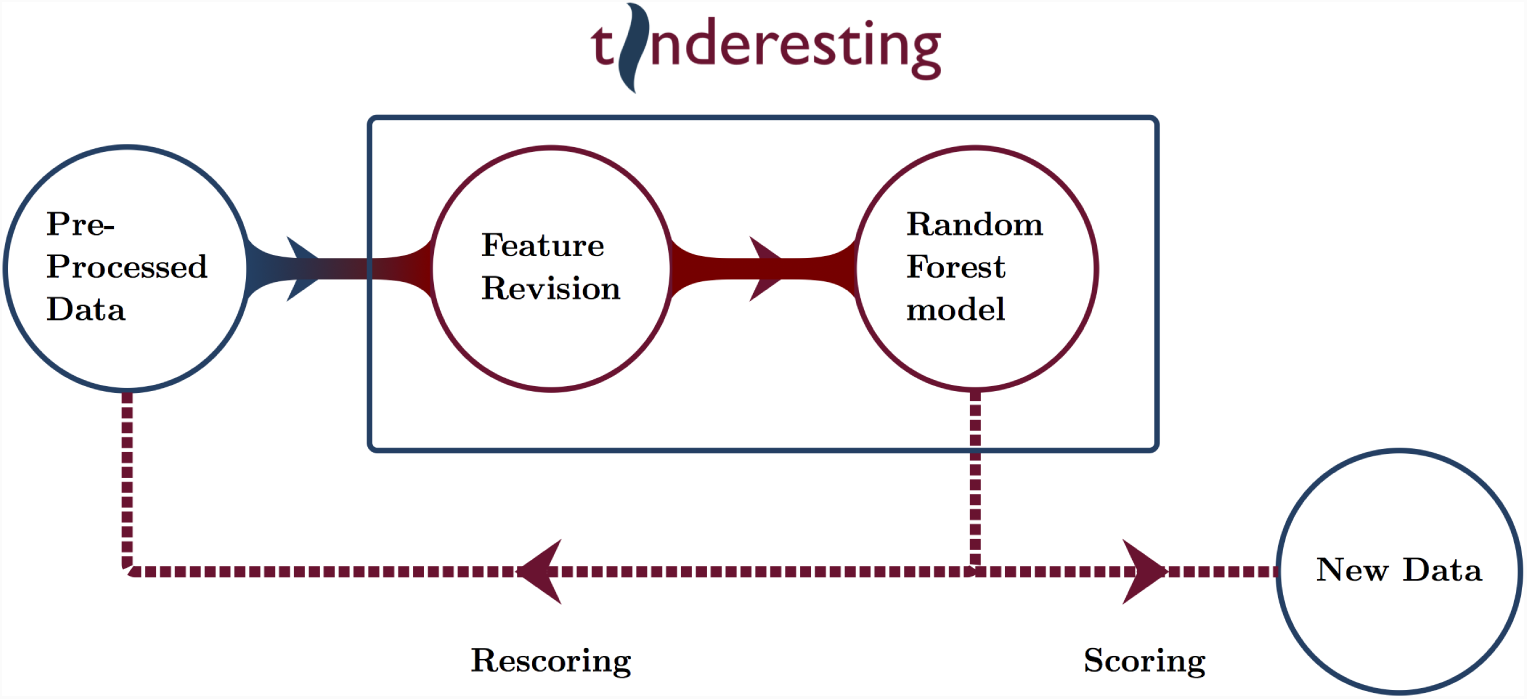
Overview of possible tinderesting workflows.

In this paper the model is used to score the dynamic metabolomics validation dataset. Performance is evaluated by using ROC and PR curves. It is important to note that, when applying tinderesting for rescoring, care needs to be taken to avoid overfitting. Specifically, the features already used to train the tinderesting model need to be removed from the dataset on which to apply rescoring.

## Results

### Simulating dynamic metabolomics data

The result of the dynamic metabolomics simulation is a time curve of the concentration for every metabolite (see Fig 4). This continuous ground truth data is sampled at discrete timepoints, corresponding to actual experiments where the underlying biological process is continuous but samples are taken at distinct time intervals because an LC-MS analysis of each sample can only be done at a limited number of time points. The resulting simulated data for a single metabolite is visualized in Fig 5. Note that the dynamics in these simulated data correspond visually to example dynamics from actual experiments [16], see Supporting information S6 for two examples.

**Fig 4.**
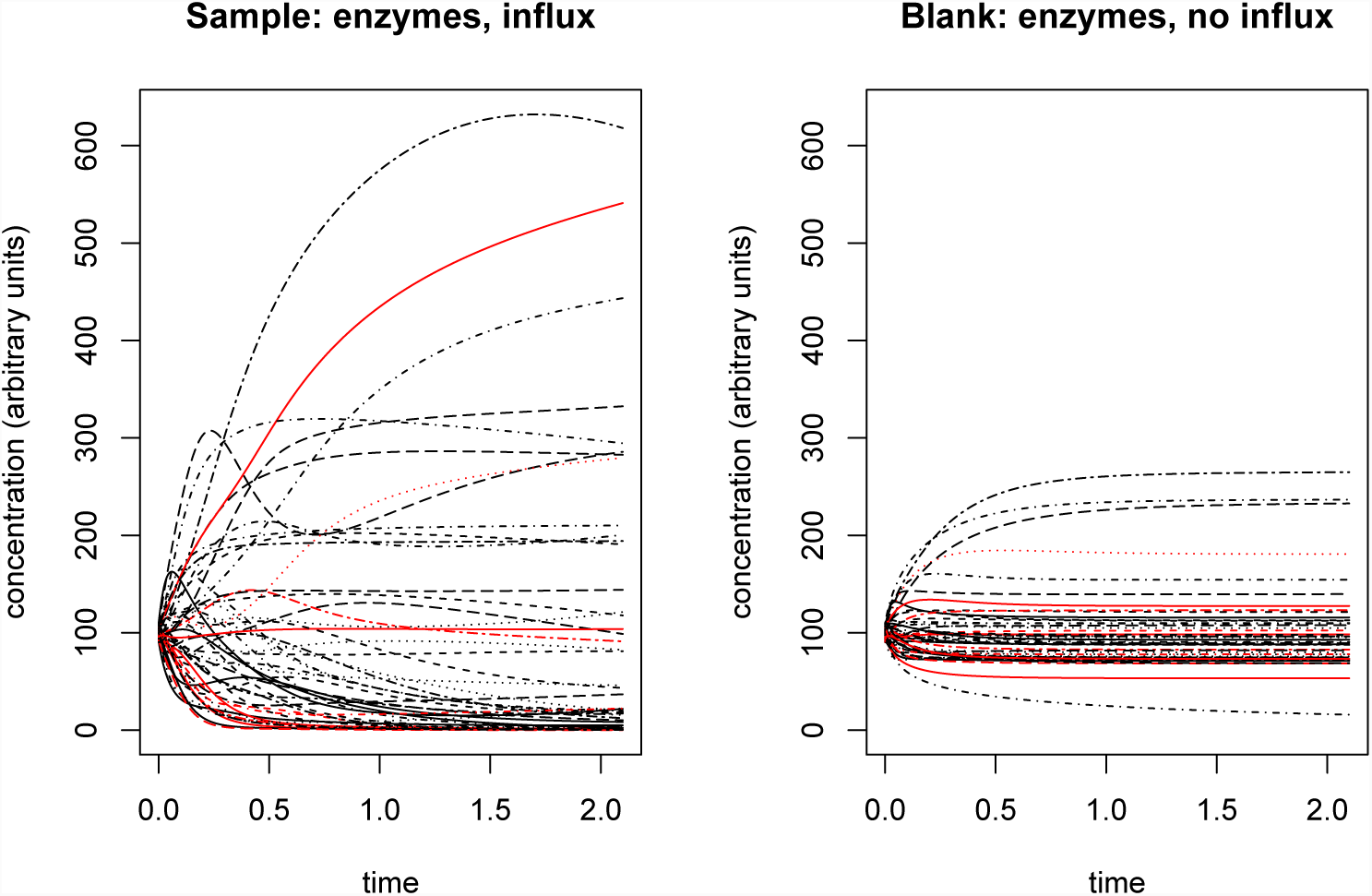
Example of time dynamics in a simulated metabolic network. Each line is a single metabolite (different line types for easier overview), a red line in the left plot indicates a metabolite (node) that receives an influx, whereas this metabolite does not receive an influx in the right plot. A single metabolic network is underlying both plots. In the left plot, the metabolic network is initiated with the same starting parameters as in the right plot (besides the random noise term). The difference between left and right is the inclusion of an external influx between time points 0.0 and 0.5 in the left plot. The difference in dynamics and end state, caused by the relatively short external influx are clear.

**Fig 5.**
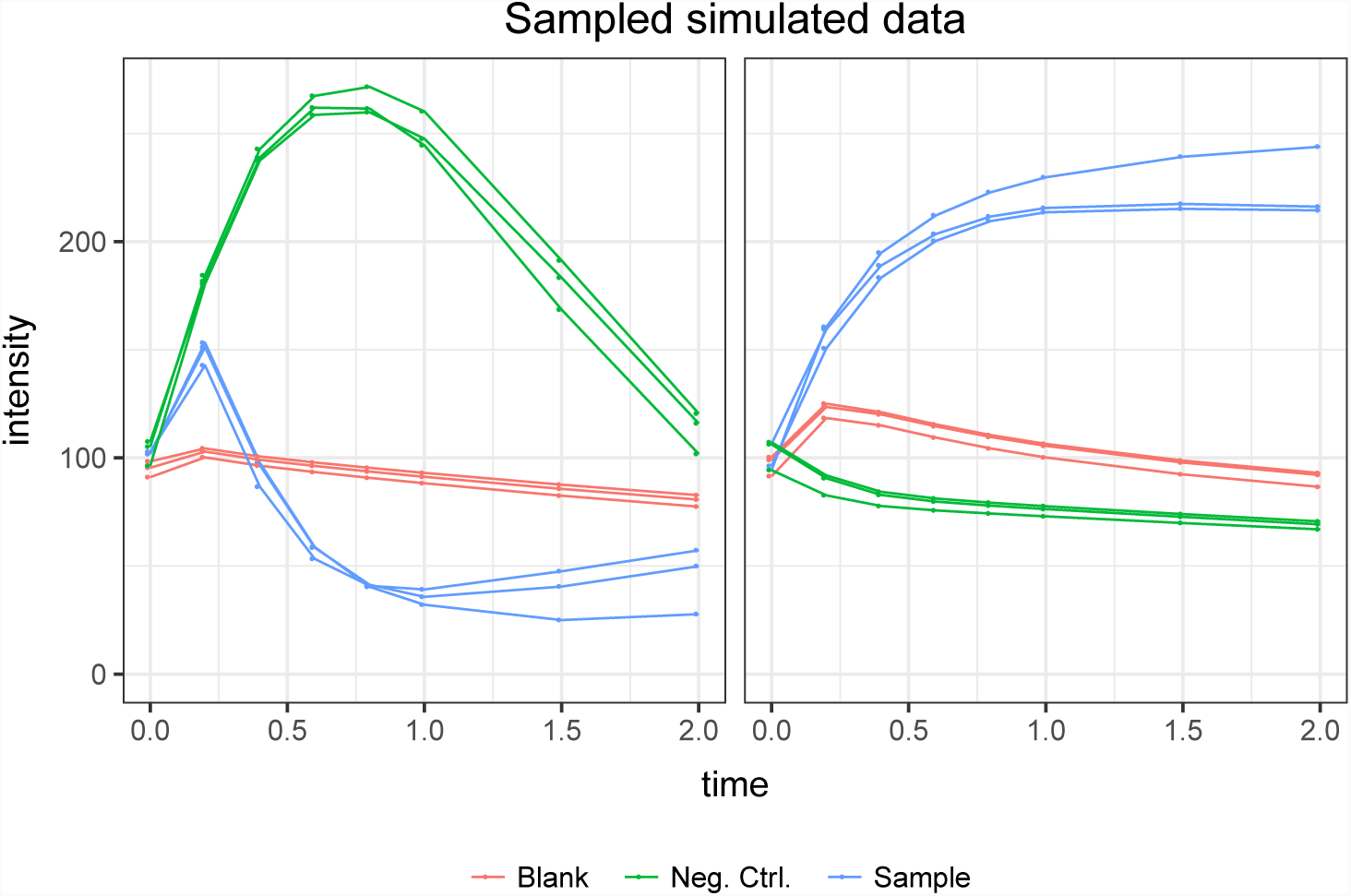
Two features (left and right) of the final simulated dataset. Taking the simulated time dynamics data from Fig 4, sampling these data and combining them by sample results in the final dataset to be used for EDGE and tinderesting. Practically, let us explain the left figure: A single green line corresponds to the sampled data of a single metabolite (line) in the left of Fig 4. That same metabolite sampled in the right half of Fig 4 results in a single red line in this image. The three lines of each class are the replicates (i.e. same network, other starting conditions)

### Statistical analysis and tinderesting

The statistical analysis is performed via the EDGE method [6, 7]. For longitudinal metabolomics, this method is robust and performant for two class experiments, i.e. case vs control. For example, the EDGE analysis for the difference between sample and blank results in an area under the ROC curve (AUC) value of 0.992 (See Fig 6 for the ROC curve), which is a near perfect classification between biologically relevant and irrelevant metabolites. Because the classes are imbalanced, the precision-recall curve is visualized in Fig 7. The same conclusions can be made. On the other hand, for the comparison between sample and negative control, the performance is significantly lower (AUC 0.675, ROC curve in Fig 6, PR curve in Fig 7), because the differences between the classes are present in all samples (albeit to a lesser degree for the 95% uninteresting metabolites). When studying the experimental setup in Fig 1, it is clear that the optimal analysis would be to only look at the difference between sample and blank. However, in actual experiments this may not be so clear and it is often the case that the experimental setup requires samples to be different from both blank and negative control samples. In this case it is non-trivial to combine the two outcomes of an EDGE analysis. Simply combining the two would lead to suboptimal results because the sample vs blank analysis should receive more weight. Setting these weights in untargeted experiments is however not straightforward. Also, by combining the analysis, the number of statistical tests per feature is effectively doubled. This theoretically leads to an increase in false positive results.

**Fig 6.**
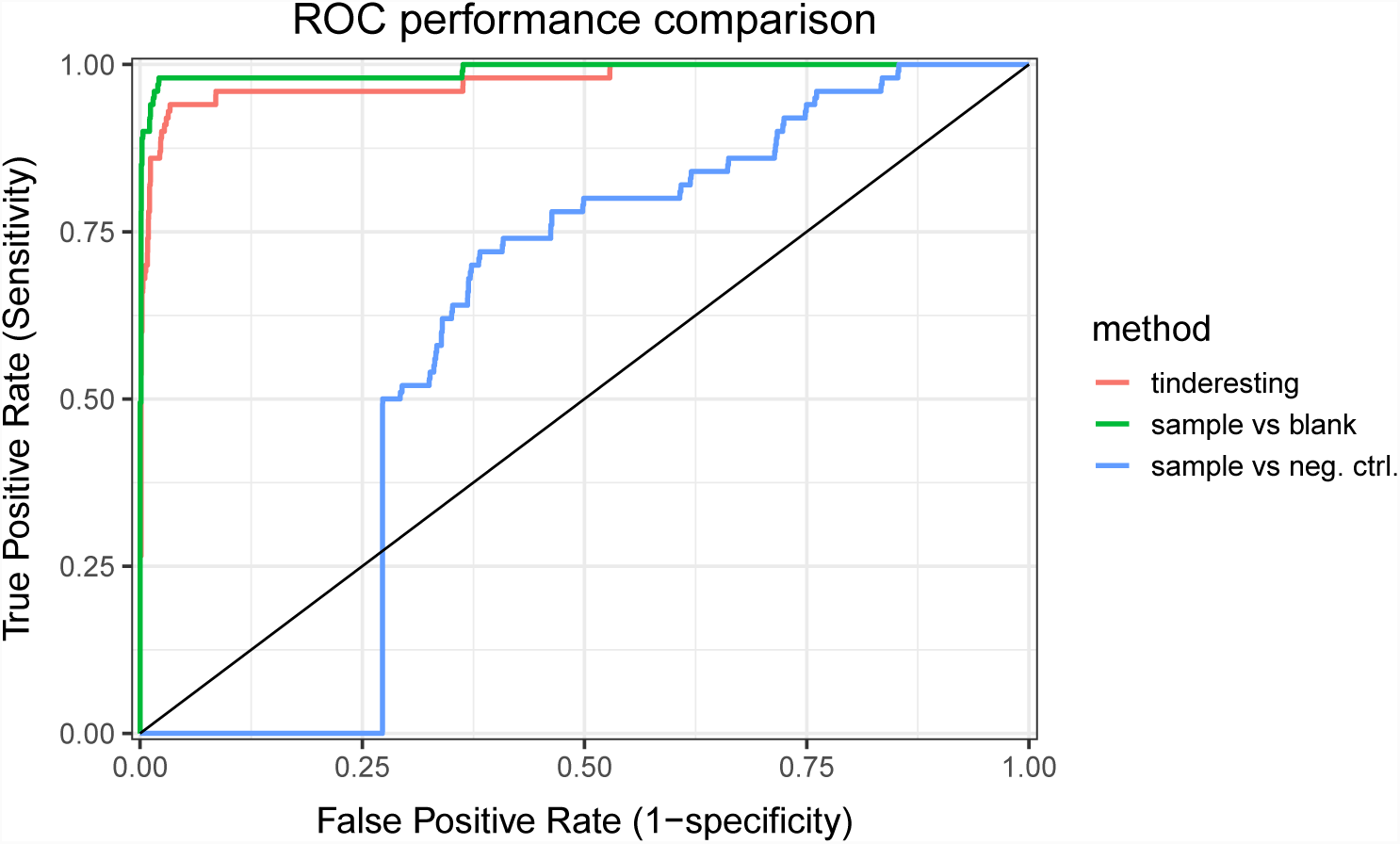
Receiver operator characteristic curves of tinderesting and EDGE. The area under the ROC curve (AUC) values are 0.977, 0.992 and 0.675 for tinderesting, sample vs blank and sample vs negative control respectively.

**Fig 7.**
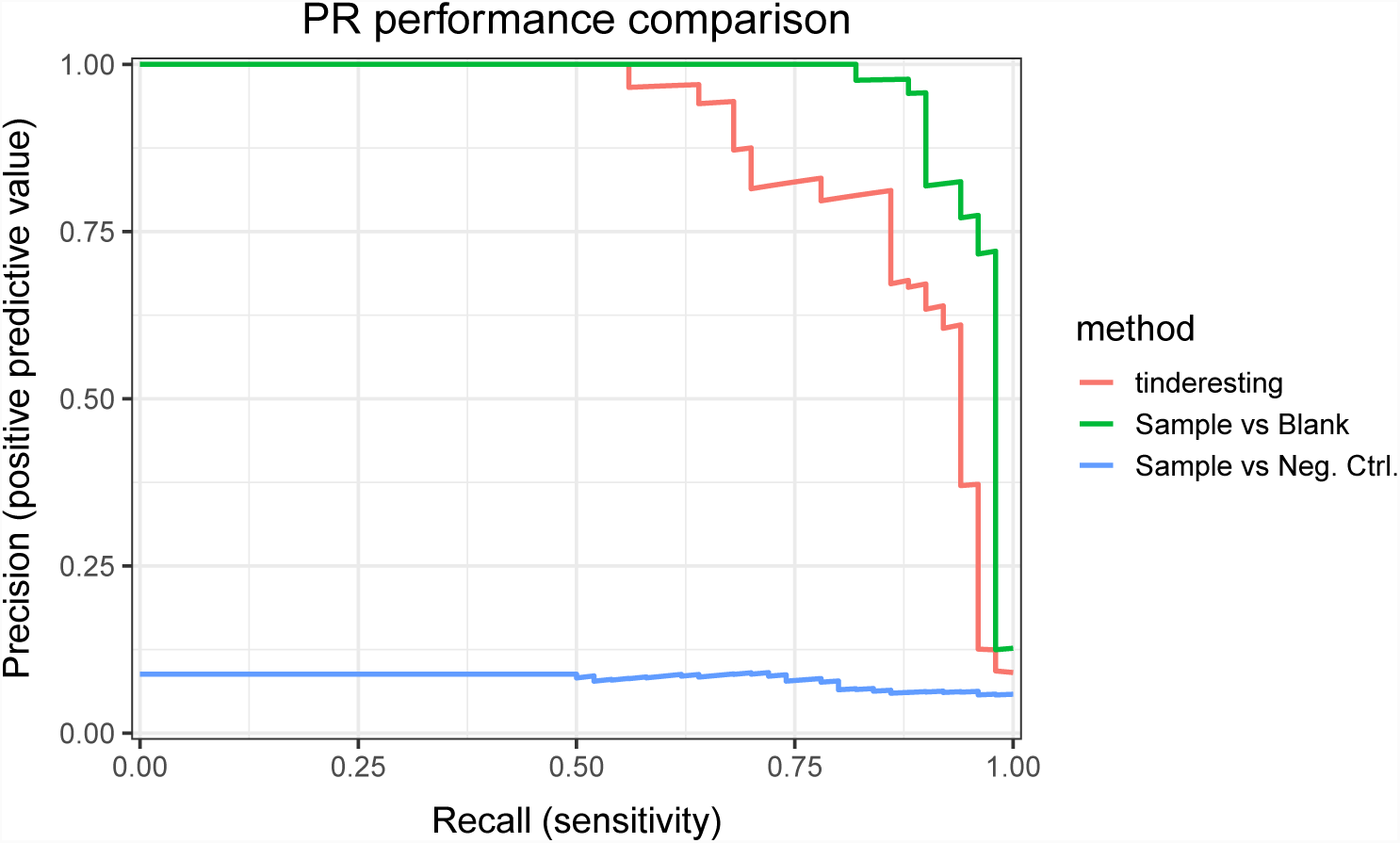
Precision-recall curves of tinderesting and EDGE. The precision-recall curves illustrate the same overall results as the ROC curves. Indicating that the performance is not due to class imbalance artefacts.

The method that is proposed in the tinderesting model circumvents these problems by training the random forest model on the data of all sample classes simultaneously. The number of classes does not have an effect on the number of tests to be performed and there are no difficulties in combining results. The outcome is a single score for each feature, independent of the number of samples, groups or classes that are present in the features. To avoid overfitting, the tinderesting model was trained on a different dataset, see Supporting information S5. The training set consisted of 185 samples: 77 interesting and 108 uninteresting are included, 15 samples for which the reviewer was uncertain where omitted. The training samples have 72 features each (8 time points per sample, 3 replicates per sample class, 3 classes). Cross validation was used to quantify the performance of the model on the training data (AUC 0.9818, see Supporting information S5 for ROC and PR curves). This model was then used on the validation dataset, specifically to predict the probabilities for samples to belong to the interesting class. The ROC curve is visualized in Fig 6 and has a corresponding AUC value of 0.977, which is only slightly below the optimal EDGE score. The precision-recall curves in Fig 7 illustrate similar results. Although there is a slight decrease in performance of tinderesting as compared to the statistical ideal situation, this is greatly offset by the advantages tinderesting offers in real life experiments with often complex designs.

An important note on imbalanced training data. In this case the training data is relatively balanced. However, if one class is greatly overrepresented compared to the other, issues may arise when training machine learning models and evaluating their performance. When the result after expert revision with tinderesting is a dataset that contains very little interesting features and many unintersting ones (or the other way around) it can be worthwhile to mitigate this situation. This can be done before the tinderesting step, for example by using EDGE upfront there can already be a first selection as to which features are possibly interesting. But also after the tinderesting revision process it is possible to account for the imbalance by using under-sampling or over-sampling. Methods such as SMOTE [8] over-sample the minority class to combat the imbalance.

Besides using performance metrics, the results can also be visualized in a different way. Fig 8 depicts a dot for each irrelevant feature (left) or relevant feature (right). The colour of this dot represents the tinderesting score and the uncorrected p-values from EDGE are plotted on the x and y positions: sample (S) vs negative control (NC) and sample (S) vs blank (B) respectively. Overall, uninteresting features have low tinderesting scores and interesting ones have high scores. Also note that the irrelevant features are spread out over the y axis as the EDGE model for sample vs negative control model underperforms overall. Most of the relevant features are plotted on top of each other making it difficult to see the actual tinderesting score. The distributions of these scores are visualized in Fig 9. The difference between relevant and irrelevant features is once again clear.

**Fig 8.**
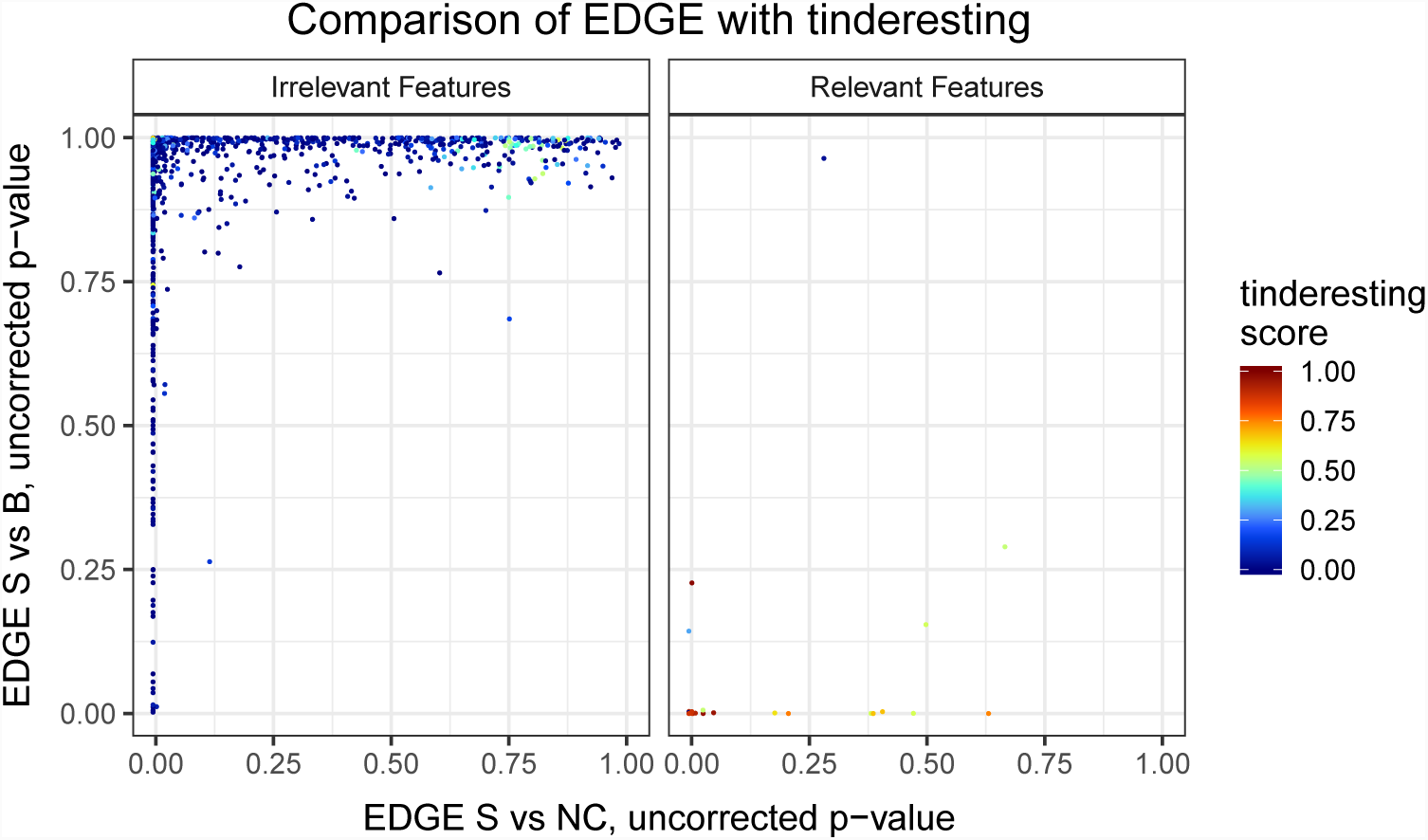
Feature scores of EDGE and tinderesting. Overall, the irrelevant features have a low tinderesting score (blue colour), and the relevant ones generally have a high score (red). The EDGE comparison of sample vs blank exhibits mostly very low values for the relevant features. The p-values of the EDGE sample vs negative control comparison are more spread out, which is consistent with the results of the performance comparison. Note that all p-values are not corrected for multiple testing. Most relevant features are plotted on top of each other in the bottom left corner on the right plot.

**Fig 9.**
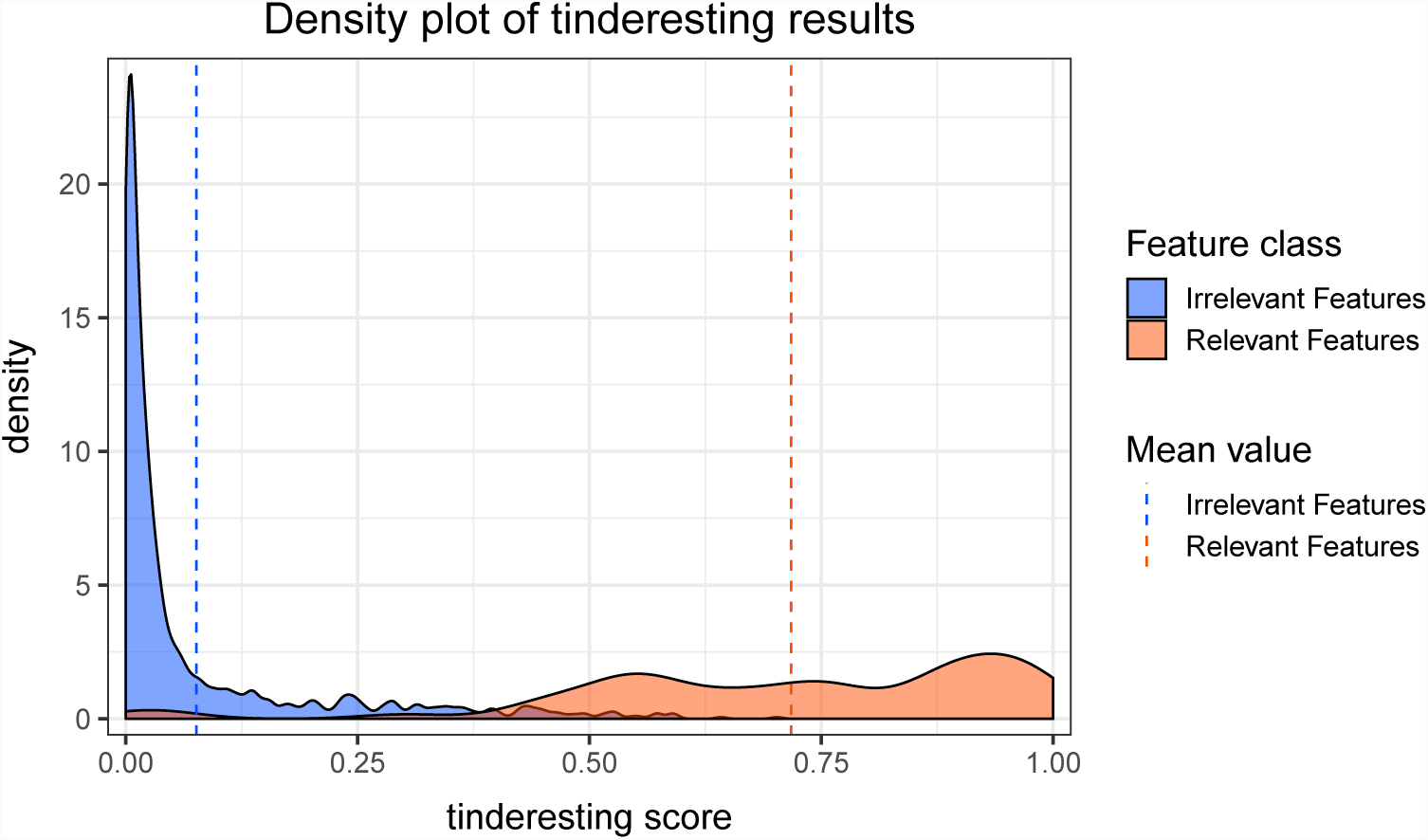
Density plot of tinderesting scores. This density plot corresponds to the ROC performance plot, there is little overlap between the distributions of irrelevant and relevant features, thus corresponding to an ROC curve with high AUC.

## Discussion & conclusions

In this paper we presented the tinderesting tool to collect expert knowledge in an easy and quick manner via a Shiny app. The expert(s) rate(s) features or images as being interesting or not. To illustrate the method we used it to analyze a simulated dynamic metabolomics dataset. Although this dataset was constructed specifically for this paper, this method of simulating data can be relevant for other tools working with longitudinal/dynamic metabolomics data as the dataset is comparable to experimental longitudinal data [16] but allows the use of a ground truth for performance evaluation (i.e. the relevant features are known).

To process this dataset with the tinderesting tool, 200 simulated features where reviewed based on their interestingness. This information was used to train a random forest model to evaluate the simulated experiment whereby 50 metabolites (features) out of 1000 are relevant. The tinderesting model performed on par with the optimal statistical setup, which is often not known in actual experiments. The merit of this approach is to be found in the agnostic nature of the method with respect to the setup of the experiment. No matter the amount of sample classes or time points in the experiment, every feature will always receive a single score. This in contrast to the case vs control statistical tools which will produce a single score (usually a p-value) per feature for every two-class comparison. A drawback on the other hand is that, the method is effectively trying to emulate the expert. If the expert is biased the model will also be biased. Hence, careful measures need to be taken. For example, the expert should review the plot of the longitudinal data of a number of features, but if these features have been linked to identified metabolites prior to expert revision, this information should be kept hidden from the reviewer. Otherwise, the reviewer might spend considerably more time reviewing the time profile of a metabolite which was upfront thought of as being interesting, thus introducing a favourable bias towards the metabolites that the researcher wants to find.

The method inherently carries the disadvantage that a new model needs to be trained for every experiment, or at least for every experiment with a different setup. If experiment A is a metabolomics study where 5 time points are measured, it is impossible to use this experiment to train a model for an otherwise identical experiment B which contains 6 time points (at least not without throwing away some information from experiment B). This is also true for experiments with different setups, different instruments, etc. The advantage, however,is that in experiments with a vast number of features, only a small part need to be reviewed for the model to work (200 for the model in this paper). This is a process that can be completed in a few minutes to maximally one hour. Also, in experiments with a standardized processing pipeline, this method of feature revision can be used to evaluate the quality of the results. A model trained to filter out false positive results from true positive results can in turn be used to find false negative results, thus effectively allowing quality control on data processing pipelines.

## Availability and future directions

The functions to generate the simulated dynamic metabolomics data are available in the MetaboLouise R package, available on github.com/Beirnaert/MetaboLouise and CRAN (awaiting approval). A light version of tinderesting is also available on the former GitHub repository. The light version uses a folder with pregenerated images for the reviewing process. This allows the easy setup of the app without the need for significant adjustments in the app to obtain the correct plots. Possible interesting additions to the workflow can come from the active learning field. When the tinderesting app, after an initial cold start phase, can query the expert with cases that lie close to the model’s decision boundary the learning rate can potentially be increased.

## Supporting information

## References

1. Giacomoni F, Le Corguillé G, Monsoor M, Landi M, Pericard P, Pétéra M, Duperier C, Tremblay-Franco M, Martin jf, Jacob D, Goulitquer S, Thévenot EA, Caron C Workflow4Metabolomics: a collaborative research infrastructure for computational metabolomics. Bioinformatics 2015; 31(9):1493–1495.

2. Weber RJM, Lawson TN, Salek RM, Ebbels TMD, Glen RC, Goodacre R, Griffin JL, Haug K, Koulman A, Moreno P, Ralser M, Steinbeck C, Dunn WB, Viant MR Computational tools and workflows in metabolomics: An international survey highlights the opportunity for harmonisation through Galaxy. Metabolomics 2017; 13(2):12.

3. Chong J, Soufan O, Li C, Caraus I, Li S, Bourque G, Wishart DS, Xia J MetaboAnalyst 4.0: towards more transparent and integrative metabolomics analysis. Nucleic Acids Research 2018; 46(W1):W486–W494.

4. Chong J, Xia J MetaboAnalystR: an R package for flexible and reproducible analysis of metabolomics data. Bioinformatics 2018

5. Smilde AK, Westerhuis JA, Hoefsloot HCJ, Bijlsma S, Rubingh CM, Vis DJ, Jellema RH, Pijl H, Roelfsema F, van der Greef J Dynamic metabolomic data analysis: a tutorial review. Metabolomics 2010; 6(1):3–17.

6. Storey JD, Xiao W, Leek JT, Tompkins RG, Davis RW Significance analysis of time course microarray experiments. PNAS 2005; 102(36):12837–12842.

7. Leek JT, Monsen E, Dabney AR, Storey JD EDGE: extraction and analysis of differential gene expression. Bioinformatics 2005; 22(4):507–508.

8. Chawla NV, Bowyer KW, Hall LO, Kegelmeyer WP SMOTE: synthetic minority over-sampling technique. Journal of artificial intelligence research 2002;16:321–57.

9. Von Ahn L, Maurer B, McMillen C, Abraham D, Blum M recaptcha: Human-based character recognition via web security measures. Science 2008; 321(5895):1465–1468.

10. Barabási AL, Albert A Emergence of scaling in random networks. Science 1999; 286(5439):509–512

11. Csardi G, Nepusz T. The igraph software package for complex network research. InterJournal, Complex Systems 2006;1695(5):1–9.

12. Pfeiffer T, Soyer OS, Bonhoeffer S. The evolution of connectivity in metabolic networks. PLoS biology 2005; 3(7):e228

13. Jeong H, Tombor B, Albert R, Oltvai ZN, Barabási AL The large-scale organization of metabolic networks. Nature 2000; 407(6804):651–654

14. Storey JD, Tibshirani R Statistical significance for genomewide studies. Proceedings of the National Academy of Sciences 2003; 100(16):9440–9445.

15. Beirnaert C, Cuyckx M, Bijttebier S MetaboMeeseeks: Helper functions for metabolomics analysis. R package version 0.1.2 https://github.com/Beirnaert/MetaboMeeseeks

16. Peeters L Manuscript in preparation

