## Supplementary material for "Using expert driven machine learning to enhance dynamic metabolomics data analysis"

### Supporting information

#### S1: Simulating biologically relevant metabolic networks

An important factor in generating random biological networks is the distribution of the connectivity. Biological networks follow a power law distribution [1$^S$], therefore it is important to maintain this property in the simulated networks. The Barabási-Albert model [2$^S$] allows the construction of networks with the appropriate distribution, see Fig 1. Note that it is important to inspect the network connectivity matrix for the following properties:

1. asymmetry: the network matrix cannot be symmetric with respect to the main diagonal, this would imply that every two metabolites that are connected, are connected in both directions. This is not the case in a biological network.

2. non-triangularity: if the network matrix exhibits a triangular shape (lower or upper triangularity), then certain nodes (metabolites) have a larger probability of having many connections. When simulating many networks this would result in, for example, metabolite A having the most connections in most of the simulations. This is non beneficial to the randomness of the generated metabolic networks. If a triangular matrix has the correct power-law distribution, the triangularity problem can be solved by randomly shuffling columns or rows.

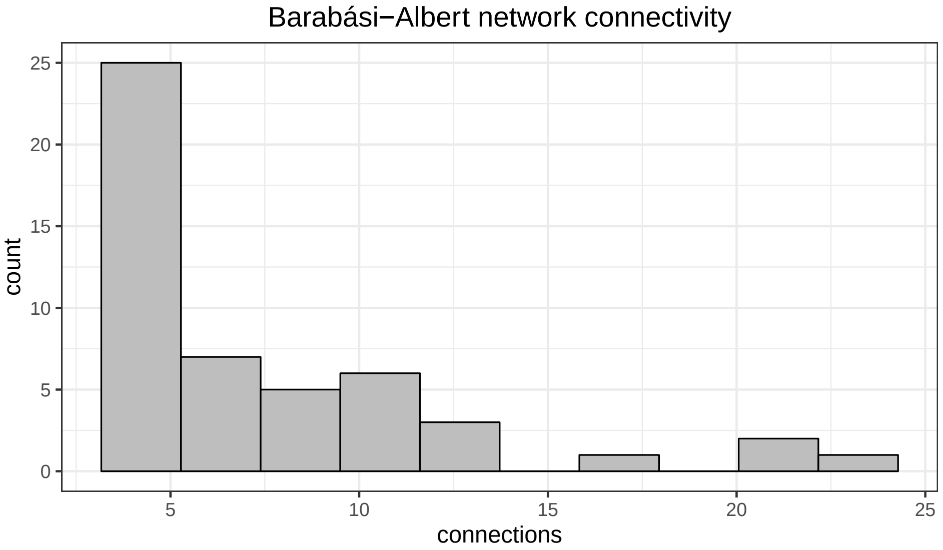


**Fig 1. Generated biological network connectivity.** The network generated with the Barabási-Albert algorithm exhibits a connectivity distribution according to a power law.

**Table 1. Simulation parameters.**

| Parameter | Value | Description |
|---|---|---|
| Nreplicates | 3 | The number of replicates for each sample. For each replicate the starting concentrations are different but the network is the same. |
| Nmetabos | 50 | The number of metabolites/nodes in the network |
| dt | 0.001 | The time step (taken sufficiently small for the Euler approximation to hold). |
| tmax | 2.1 | The end time. |
| start_concentration | 100 | The starting concentration of each metabolite without noise. |
| max_abs_concentration_noise | 10 | The maximal absolute value of the noise that is added to starting concentration. Set at 1/10th of start_concentration. |
| influx_tmax | 0.5 | The time point at which the influx (if present) stops. |
| N_influx | 10 | The number of metabolites that receive an influx. |
| Neg_ctrl_protein_factor_strong | 0.01 | The factor by which the rates are multiplied in the case of a "no influx, no enzymes" simulation (see Fig 4). |
| Neg_ctrl_protein_factor_weak | 0.5 | The factor by which the rates are multiplied in the case of a "no influx, little enzymes" simulation (see Figs 4). |
| Neg_ctrl_protein_fraction | 0.5 | The fraction of enzymes that receive the strong negative control protein factor. |
| BApower | 0.5 | Parameter of the Barabási-Albert graph model generator from the igraph R package [3S]: The power of the preferential attachment |
| BA_mValue | 4 | Parameter of the Barabási-Albert graph model generator from the igraph R package: the number of edges to add in each time step |

#### S2: Dynamics of a metabolic network, the mathematics of change over time

For our toy example the following small network [4S] can be used:

The nodes represent metabolites, A, B, C, etc. that have certain concentrations denoted as $A$, $B$, $C$ respectively. These concentrations evolve based on the enzymes x, y and z. These are effectively the rates that govern the flow and are denoted as $x$, $y$ and $z$ respectively. Thus, the evolution of the concentration over time can be written according to the following ordinary differential equations:

$$\frac{\mathrm{d}A}{\mathrm{d}t} = -(x+y)A$$
$$\frac{\mathrm{d}B}{\mathrm{d}t} = xA$$
$$\frac{\mathrm{d}C}{\mathrm{d}t} = yA - (x+z)C$$
$$\vdots$$

To numerically solve these equations the Euler approximation is used by substituting the following finite difference scheme into the equations for the metabolic network

$$\frac{\mathrm{d}f}{\mathrm{d}t} \approx \frac{f(t+\delta t) - f(t)}{\delta t}$$

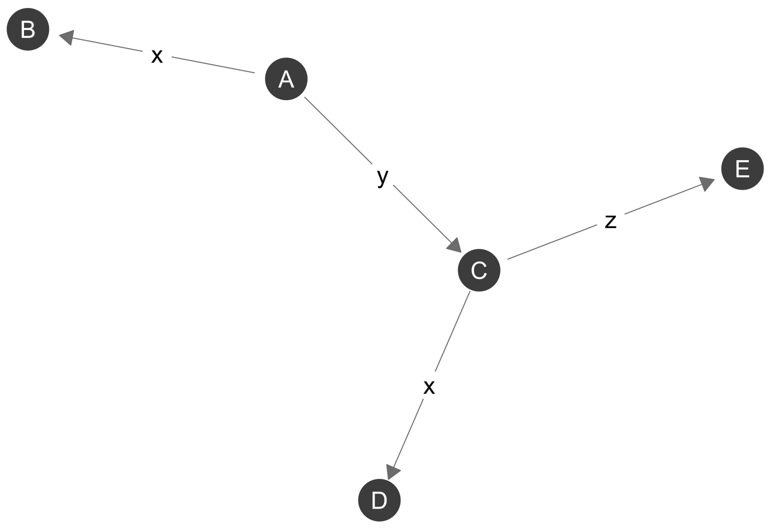


**Fig 2. Metabolomic network example.** Nodes are metabolites and the edges represent flow between metabolites. These flows are rates based on concentrations of enzymes.

by writing $f(t+\delta t)$ in the following shorthand notation $f_{t+\delta t}$ we obtain the following

$$\begin{aligned}\frac{A_{t+\delta t}-A_t}{\delta t} &= -(x+y)A_t\\ \frac{B_{t+\delta t}-B_t}{\delta t} &= xA_t\\ \frac{C_{t+\delta t}-C_t}{\delta t} &= yA_t-(x+z)C_t\\ &\vdots\end{aligned}$$

Reworking this gives the final result for the state of the network at time point $t+\delta t$ in function of the state at time point $t$

$$\begin{aligned}A_{t+\delta t} &= -\delta t(x+y)A_t + A_t\\ B_{t+\delta t} &= \delta t x A_t + B_t\\ C_{t+\delta t} &= \delta t(yA_t-(x+z)C_t)+C_t\\ &\vdots\end{aligned}$$

When choosing $\delta t$ sufficiently small the Euler approximation is valid. As mentioned in the manuscript, the rates are influenced by the concentration of metabolites. Specifically, when the concentration of a metabolite increases, the rates that deplete that metabolite will also increase, up to a certain maximum. This maximum has been arbitrarily set at twice the starting rate. For example, the equation governing the evolution of rate y is

$$y(A_t) = \frac{y^{max}}{1+\exp[-k(A_t-A_0)]}$$

#### S3: Multiple testing correction problem for EDGE and metabolomics

A critical note on p-values and multiple testing correction has to be made. The method performs a test for an improvement in model performance. This is effectively a one-sided test (The model with two curves is better than the model with one curve). Large numbers of these one-sided tests can result in a bimodal p-value distribution. Such a bimodal p-value distribution can prevent the adequate application of the often used FDR correction, as it uses the distribution of the p-values to estimate the $\pi_0$ value (proportion of true nulls), and this distribution is assumed to be roughly uniform in the high p-value region [$6^S$]. With regard to bimodal p-values, such a distribution can further be caused by a large number of features without any difference between sample classes, This can often occur in biological experiments as large numbers of (uninformative) features are measured. In this case, the bimodal distribution problem can be solved by either removing these features or by applying surrogate variable analysis techniques, which can also be used to remove batch effects and other unwanted variation [$7^S$, $8^S$].

#### S4: Shiny app and user interaction

To review the quality of the significant features and to train the machine learning model, we constructed a Shiny web app called tinderesting. Shiny is an R package to produce such interactive web apps that require little effort to construct and can easily be run on a server. The app queries the user for the quality of features, and uses these results to build a model in the background. See Fig 3. All responses are stored in an SQLite database for straightforward storage and access.

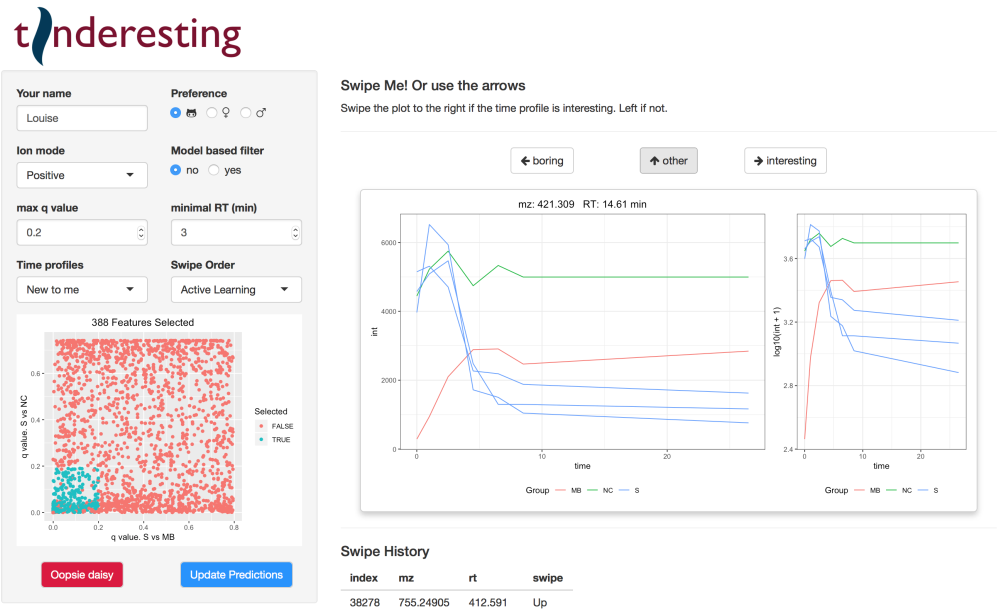


**Fig 3. Visualization of the full tinderesting app.** The user selects the subset of the data on the left hand side, the selected subset is visualized. Next, features appear on the right hand side that need reviewing by the expert. The results are logged at the bottom of the app and simultaneously stored in an SQL database.

#### S5: Performance comparison of machine learning models.

To find the optimal machine learning model for the longitudinal metabolomics data the predictive performance was evaluated on an other dataset than the one discussed in the paper. The setup is visualized in Fig 4, the seed is set differently so as to obtain different networks (to avoid overfitting) than those of the experiment in the paper. The models are

1. a random forest with 500 trees, 8 variables sampled at each split, minimum size of terminal node 1, sampling with replacement (the default parameters from the randomForest package [$9^S$]),
2. a support vector machine with a radial basis kernel (rest of parameters are default from e1071 R package, [$10^S$])
3. a naive Bayes classifier (default e1071 R package [$10^S$] parameters, no priors set).

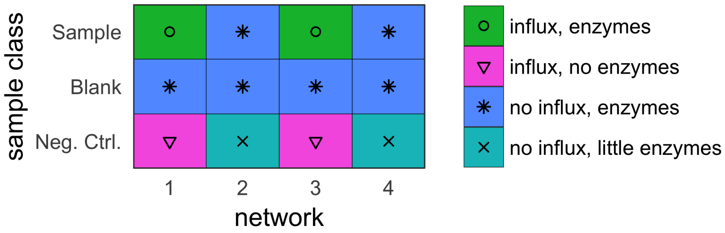


**Fig 4. External tinderesting data, simulated setup.**

The performance of the three classifiers was compared in a 10 fold cross validation setup, each repeated 20 times with different folds (to get an estimate of the ROC curve variability). The results are visualized in Fig 5 where it is clearly shown that the random forest model consistently outperforms the others. The code to run these models and obtain similar plots is available in the MetaboMeeseeks R package [$11^S$]. For completeness, the Precison-Recall plot of the Random forest model is plotted in Fig 6. The random forest model trained with this simulated data is used to compare to the EDGE model. This is to avoid overfitting on the data and to allow a fair comparison between EDGE and tinderesting.

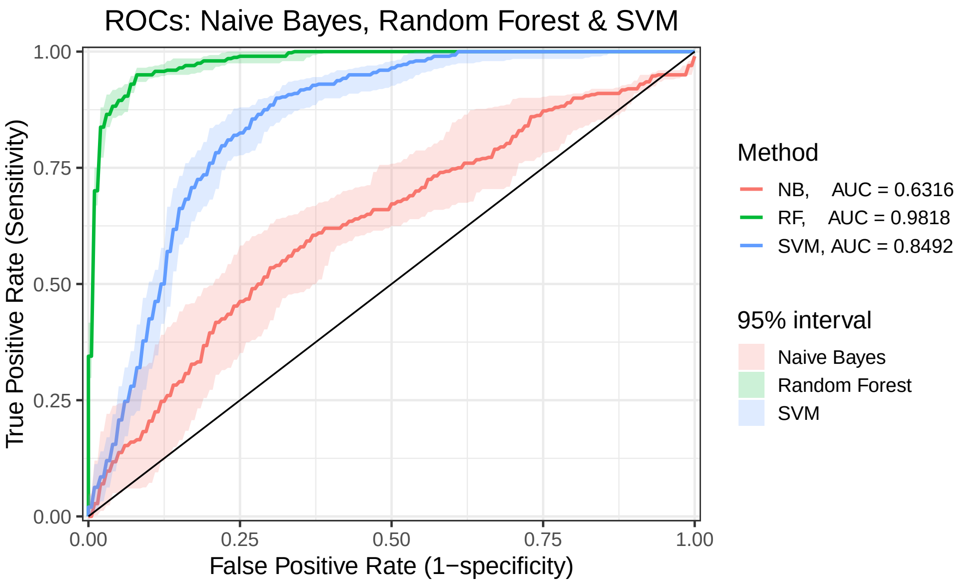


**Fig 5. ROC curves for random forest, SVM and naive Bayes classifiers.** Classifier performance is represented with the respective receiver-operator-characteristics. Comparing the area under the ROC curves (AUC values) justifies the choice for the random forest classifier.

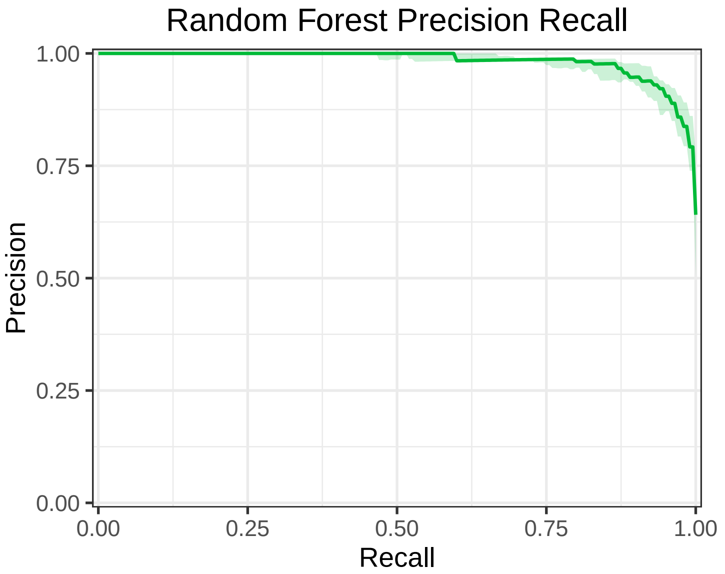


**Fig 6. Precison-recall curve for the random forest classifier.**

#### S6: Examples of actual dynamic metabolomics data

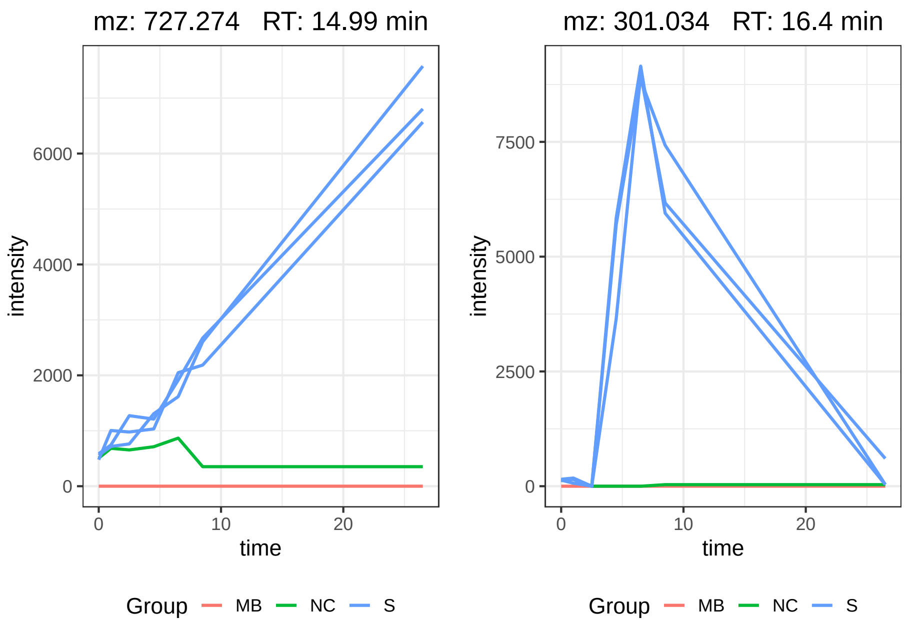


**Fig 7. Two examples from a dynamic metabolomics experiment.** Visualization of two features from the processed LC-MS dataset [5S]. The sample class (S) shows different behavior over time compared to the other classes (MB = method blank, NC = negative control). Left, a compound is formed in the sample class and right, a compound is formed and broken down again.

#### Supporting references


1S. Jeong H, Tombor B, Albert R, Oltvai ZN, Barabási AL The large-scale organization of metabolic networks. Nature 2000; 407(6804):651–654

2S. Barabási AL, Albert A Emergence of scaling in random networks. Science 1999; 286(5439):509-512

3S. Csardi G, Nepusz T. The igraph software package for complex network research. InterJournal, Complex Systems 2006;1695(5):1-9.

4S. Tyner S, Briatte F, Hofmann H. Network Visualization with ggplot2. The R Journal 2017;

5S. Peeters L Manuscript in preparation

6S. Storey JD, Tibshirani R Statistical significance for genomewide studies. Proceedings of the National Academy of Sciences 2003; 100(16):9440–9445.

7S. Leek JT, Storey JD Capturing Heterogeneity in Gene Expression Studies by Surrogate Variable Analysis. PLoS Genetics 2007; 3(9):e161

8S. Leek JT, Johnson WE, Parker HS, Jaffe AE, Storey JD. The sva package for removing batch effects and other unwanted variation in high-throughput experiments. Bioinformatics 2012; 28(6):882-3.

9[S]. Liaw A, Wiener M Classification and Regression by randomForest. R News 2002; 2(3): 18–22.

10[S]. Meyer D, Dimitriadou E, Hornik K, Weingessel A, Leisch F e1071: Misc Functions of the Department of Statistics, Probability Theory Group (Formerly: E1071), TU Wien. R package version 1.7-0. https://CRAN.R-project.org/package=e1071

11[S]. Beirnaert C, Cuyckx M, Bijttebier S MetaboMeeseeks: Helper functions for metabolomics analysis. R package version 0.1.2 https://github.com/Beirnaert/MetaboMeeseeks
